# Humidity Controls the Timing and Persistence of Ozone Injury in Citrus: Linking Leaf Physiology and Regional Canopy Responses

**DOI:** 10.1101/2025.10.28.685042

**Authors:** Luka Mamić, Mj Riches, Delphine K. Farmer

## Abstract

Tropospheric ozone (O_3_) is a major air pollutant that threatens crop productivity, yet its effects depend strongly on environmental conditions that regulate plant O_3_ uptake. Here, we explore how citrus, an O_3_-sensitive perennial crop, responds to O_3_ exposure under humid subtropical (Florida) and semi-arid (California) climates. In controlled chamber experiments, Meyer *lemon* trees exposed to moderate O_3_ concentrations (80 ppb for 4 h d^−1^ over four days) showed a faster and more persistent decline in the maximum photosystem II efficiency (F_v_/F_m_) under humid air, while under dry air the response was delayed by one day and reversible. This humidity-dependent behavior reflects differences in stomatal conductance (g_s_) where high humidity maintains open stomata and accelerates O_3_ flux and dry air limits uptake but enhances slower non-stomatal injury pathways.

At regional scale, satellite solar-induced chlorophyll fluorescence (SIF) from Sentinel-5P TROPOMI revealed similar patterns. In Florida, SIF decreased significantly during O_3_-episode weeks and remained low for up to three weeks, while in California it showed a brief rebound before a delayed decline – mirroring the timing observed in the chamber experiment. Analysis of the SIF and gross primary productivity (GPP) relationship further showed that O_3_ decoupled canopy fluorescence from productivity in the dry region, whereas drought stress weakened this coupling in the humid region, indicating a climate-specific shift in the dominant stressor.

We demonstrate and argue that humidity governs both the timing and persistence of O_3_ injury, linking leaf-level physiology to regional canopy responses. These findings emphasize that effective O_3_-risk assessments for perennial crops must incorporate local humidity and vapor pressure deficit conditions and both stomatal and non-stomatal deposition pathways.

## 1. Introduction

Air pollution is a significant threat to vegetation, with ground-level ozone (O_3_) as one of the most established pollutants known to damage plants (Ashmore, 2005; Emberson, 2020; Emberson et al., 2009; He et al., 2021; Manono et al., 2025). However, plants are one of the most important sinks for tropospheric O_3_, thus potentially improving air quality (Clifton et al., 2020; Fares et al., 2014). This dual role is particularly relevant for perennials such as citrus, where both stomatal and non-stomatal processes contribute significantly to O_3_ uptake and surface removal (Fares et al., 2010).

O_3_ exposure disrupts physiological processes in plants – such as photosynthesis and chlorophyll fluorescence – leading to reduced crop yields (Mamić et al., 2025a; Mills et al., 2007; Tai et al., 2014). In citrus, prolonged exposure to elevated O_3_ (≥ 80-120 ppb) has been shown to lower chlorophyll and carotenoid concentrations, decrease the maximum quantum efficiency of photosystem II (F_v_/F_m_), and induce oxidative stress through lipid peroxidation and ascorbate depletion (Calatayud et al., 2006; Delgado-Saborit & Esteve-Cano, 2008).

The physiological responses of plants vary under different environmental conditions, further complicating the interaction between O_3_ and vegetation (Martin et al., 2025; Sun et al., 2022) – especially due to the different stomatal behavior (Clifton et al., 2020). Stomatal conductance (g_s_) is driven by environmental factors such as moisture availability (Medlyn et al., 2011). In citrus, g_s_ largely determines O_3_ flux into the leaf, but non-stomatal deposition via cuticular surfaces and reactions with biogenic volatile organic compounds can also become relevant under high O_3_ or water-limited conditions (Fares et al., 2010). These processes determine not only the magnitude but also the timing and persistence of O_3_ injury, as shown in recent deposition studies (Sun et al., 2022; Wong et al., 2022).

In humid climates, plants tend to open their stomata to maximize carbon dioxide (CO_2_) intake, while in dry environments they often partially or fully close their stomata to conserve water (Wong et al., 2022). These contrasting behaviors suggest that identical ambient O_3_ exposure could have different effects on plant physiological stress in a humid subtropical versus a hot semi-arid environment. However, early experimental studies shows that high relative humidity (RH) can enhance O_3_ injury by keeping stomata open (Otto & Daines, 1969), whereas under dry air, O_3_ uptake declines but oxidative stress may still occur within intercellular spaces due to internal accumulation (Fares et al., 2010). Some studies indicate that drought-induced stomatal closure might not entirely shield plants from O_3_ damage; O_3_ effects (e.g. on photosynthetic capacity and yield) can persist even under water-limited conditions (Martin et al., 2025; Wong et al., 2022). In citrus, chronic O_3_ exposure has been associated with reduced F_v_/F_m_ and impaired photosynthetic recovery even when moderate drought limits g_s_ (Delgado-Saborit & Esteve-Cano, 2008). In soybean field experiments, elevated O_3_ was found to reduce photosynthetic capacity and yield additively to drought stress, without preventing normal stomatal closure by drought (Martin et al., 2025). Thus, the question remains: Do plants respond differently to a given O_3_ exposure under humid versus arid conditions, and how does atmospheric humidity (or drought) modulate O_3_’s impact on plant physiological function?

To address this knowledge gap, we conducted an integrative study combining controlled experiments and regional remote sensing observations. We chose citrus plants because citrus is both economically important (Liu et al., 2012) and grown in regions with contrasting climates – notably Florida’s humid subtropical climate and California’s semi-arid climate. O_3_ pollution is a concern in both areas (both experience similar ambient O_3_ levels on the order of ∼40 ppb on a weekly average basis year-round), yet humidity and water availability differ greatly. In the first part of this study, we performed O_3_ fumigation chamber experiments on potted Meyer lemon (*Citrus × meyeri*) to quantify leaf-level physiological responses to O_3_ under controlled humid versus dry conditions. We focused on the F_v_/F_m_ as a sensitive indicator of photosynthetic performance and oxidative stress, building onto previous citrus-O_3_ response studies (Calatayud et al., 2006; Delgado-Saborit & Esteve-Cano, 2008). In the second part, we extended our analysis by examining satellite-derived plant health indicators in two key citrus-growing regions: the Indian River County in Florida (humid subtropical) and the Fresno County, California (hot semi-arid). We analyzed satellite measurements of solar-induced chlorophyll fluorescence (SIF) from the Sentinel-5P TROPOMI instrument, which provides a proxy for canopy photosynthesis (Guanter et al., 2021), evaluating its relationship to gross primary production (GPP) and accumulated O_3_ exposure above 40 ppb (AOT40) across citrus regions. By integrating these approaches, we link leaf-level O_3_ injury mechanisms to regional canopy productivity, testing how humidity and drought modulate O_3_ stress from physiology to landscape.

## 2. Methodology

### 2.1. Plant-level chamber experiments

To isolate the effect of humidity from O_3_, we conducted controlled fumigation experiments on young citrus trees under alternating humidity regimes within a single fumigation chamber. Meyer lemon trees were grown in pots and acclimated to laboratory conditions before starting with the experiment.

All environmental parameters – temperature, light, and CO2 concentration – were kept constant, while RH varied to simulate contrasting climate conditions: ∼25% RH (dry, semi-arid) and ∼75% RH (humid, subtropical). Air temperature was maintained at 25 ± 2 °C, light intensity at 1000 µmol m^−2^ s^−1^, and CO2 mixing ratios at ∼400 ppm. This ensured that the only planned environmental difference between treatment groups was the RH of the air.

Plants were exposed to 80 ± 10 ppb O_3_ for four hours per day over four consecutive days. O_3_ was generated from a pen-ray lamp (Analytik Jena AG) connected to the chamber’s dry air inlet. Between fumigation periods, plants were maintained at ambient laboratory O_3_ levels (∼30 ppb daily mean).

To assess photosynthetic efficiency and potential photoinhibition, maximum quantum yield of photosystem II (F_v_/F_m_) was measured each day on five leaves per plant using a Li-Cor LI-6800 portable photosynthesis system (PPS). For dark-adapted F_v_/F_m_ measurements, leaves were covered with aluminum foil for at least one hour prior to measurement to allow full relaxation of photosystem II reaction centers (Baker, 2008; Jia et al., 2019; Kobayakawa et al., 2013; Maxwell & Johnson, 2000).

Measurements were taken before (pre-treatment) and after (post-treatment) each O_3_ exposure. Control plants underwent the same procedure, but without O_3_ fumigation. Three replicate plants were used per condition: dry/O_3_, humid/O_3_, and controls (one dry and two humid). Leaf-level data were averaged per plant for analysis.

Although all plants were well-watered, the contrasting RH conditions produced distinct vapor pressure deficits (VPDs) consequently controlling stomatal behavior and O_3_ uptake (Wang et al., 2009). Including control plants under both humidity regimes allowed separation of humidity-driven effects from O_3_-specific responses.

Given the limited sample size and the need to distinguish biological from measurement outliers (Julián et al., 2024) we applied the robust method of Leys et al. (2013), excluding values beyond three median absolute deviations. This minimized the influence of anomalous points while keeping representative physiological responses.

Additional details on the chamber setup are provided in Mamić et al. (2025a). We note that chamber and satellite data are not directly comparable due to their large difference in spatial scales (leaf versus canopy). Rather, our objective was to test whether the physiological O_3_ stress observed under controlled humidity conditions is consistent with patterns detected at the regional scale. Results from plant-level chamber experiments are presented and discussed in section 3.1.

### 2.2 . Regional remote sensing analysis

To see if the leaf-level mechanisms observed in the chamber experiments also appear at the canopy scale, we analyzed satellite remote sensing data for two major U.S. citrus-growing regions with contrasting climates: Fresno County, California (hot semi-arid) and Indian River County, Florida (humid subtropical) (see Section 3.2). Both regions have comparable mean ambient O_3_ levels (∼35-45 ppb annually) but differ strongly in RH, VPD, and drought frequency, providing a natural contrast in environmental control of O_3_ uptake (**Fig S1**).

Although citrus is an evergreen crop, its photosynthetic activity follows a clear annual cycle, as visible in satellite SIF and GPP time series (Mamić et al., 2025b). To focus on periods of active canopy photosynthesis, we selected weeks corresponding to the regional photosynthetic seasons defined in Mamić et al. (2025b) using normalized SIF signatures (see https://lukmam.shinyapps.io/sifsignaturesapp/). In the humid subtropical climate of Florida, citrus exhibits high photosynthetic efficiency from April to October (weeks 15-41), whereas in the hot semi-arid climate of California, activity peaks from February to August (weeks 8-35), with generally lower SIF amplitudes. Restricting the analysis to these periods ensured that observed variability reflected physiological responses rather than seasonal dormancy.

For each region, we defined a few areas of interest covering most citrus orchards based on United States Department of Agriculture land cover maps. To capture canopy-level photosynthetic activity, we used SIF derived from Sentinel-5P. The ESA NOVELTIS TROPOSIF dataset provides corrected baseline SIF Level-2 product (far-red 743-758 nm; mW m^-2^ sr^-1^ nm^-1^) at a spatial resolution of 3.5 × 7 km (available at https://s5p-troposif.noveltis.fr/data-access/) (Guanter et al., 2021). We analyzed data from January 2019 to December 2021. Negative and extreme SIF values were removed, and remaining observations were spatially averaged within each area of interest. Daily data were aggregated to weekly means to minimize retrieval noise and atmospheric variability.

For comparison, GPP data was obtained from the MODIS model (8-day composite; 250 m spatial resolution; kg C m^−2^ 8-day^−1^) (Robinson et al., 2018). GPP was divided by 8 to get approximate daily values and then summed across the week to match the aggregated SIF resolution. These paired datasets enabled evaluation of SIF-GPP coupling as an indicator of canopy photosynthetic efficiency under varying stress conditions (section 3.3).

Meteorological variables – VPD and RH – were obtained from the GRIDMET model (4-km resolution) (Abatzoglou, 2013). Weekly Drought Severity and Coverage Index (DSCI) values were downloaded from the U.S. Drought Monitor (https://droughtmonitor.unl.edu/) to quantify drought intensity through time.

Hourly ground-level O_3_ concentrations were obtained from an EPA Air Quality System monitoring stations in Fresno and Indian River Lagoon (https://aqs.epa.gov/aqsweb/airdata/download_files.html#Raw). The accumulated O_3_ exposure above 40 ppb (AOT40) was calculated as the sum of O_3_ exceeding 40 ppb during daylight hours (08:00–20:00) within each week. Although mean weekly O_3_ levels are similar between regions, Fresno exhibits notably higher AOT40, indicating more frequent high-O_3_ events.

Because plants are adapted to local ambient O_3_, we defined O_3_ episodes as statistically extreme deviations from this baseline. Therefore, weeks in which AOT40 exceeded the regional, year-specific median by more than three median absolute deviations were labeled as O_3_ episodes. To minimize drought confounding, AOT40 was residualized against DSCI using a linear model (AOT40 ∼ DSCI), and episode classification was based on the residuals (drought-corrected O_3_ anomalies). Weeks within ± 3 weeks of any episode were excluded from the non-episode reference group to ensure clean separation between high-O_3_ and background periods. The resulting isolated O_3_ episodes for each region and year are shown in **Fig. S2**.

Weekly SIF, GPP, and drought data were integrated to evaluate canopy photosynthetic responses to O_3_ stress under contrasting humidity regimes during each region’s active season. Depressions in SIF or declines in the SIF-GPP relationship during high-O_3_ weeks were interpreted as evidence of O_3_-induced down-regulation of photosynthesis. This regional analysis complements the controlled chamber experiments by testing whether the humidity-dependent timing and persistence of O_3_ injury observed at the leaf level (section 3.1) are expressed at the canopy scale (section 3.2) and how they influence productivity coupling (section 3.3) across real citrus landscapes. By integrating mechanistic and satellite observations, this multi-scale framework links O_3_ stress physiology with ecosystem-level carbon dynamics, providing insight into how environmental context determines O_3_ impacts on perennial crops.

## 3. Results and Discussion

### 3.1. Leaf-level F_v_/F_m_ responses to repeated O_3_ exposure under contrasting humidity – stomatal versus non-stomatal control of injury timing and persistence

Across four consecutive exposure days (80 ppb O_3_ for 4 h d^−1^), the F_v_/F_m_ diverged clearly between humidity treatments (**Fig. 1**). Under humid air (∼75% RH), F_v_/F_m_ in O_3_-fumigated plants tracked the controls through the first two days, but separated on the third day of exposure – the first day in which we noticed clear differences between non-fumigated and fumigated plants, which diverged even more on the fourth day (**Fig. 1a**). After two days without elevated O_3_ chamber exposure (**Fig. 1b**), F_v_/F_m_ recovery remained incomplete and significantly lower than controls (p = 0.002).

**Fig. 1.**
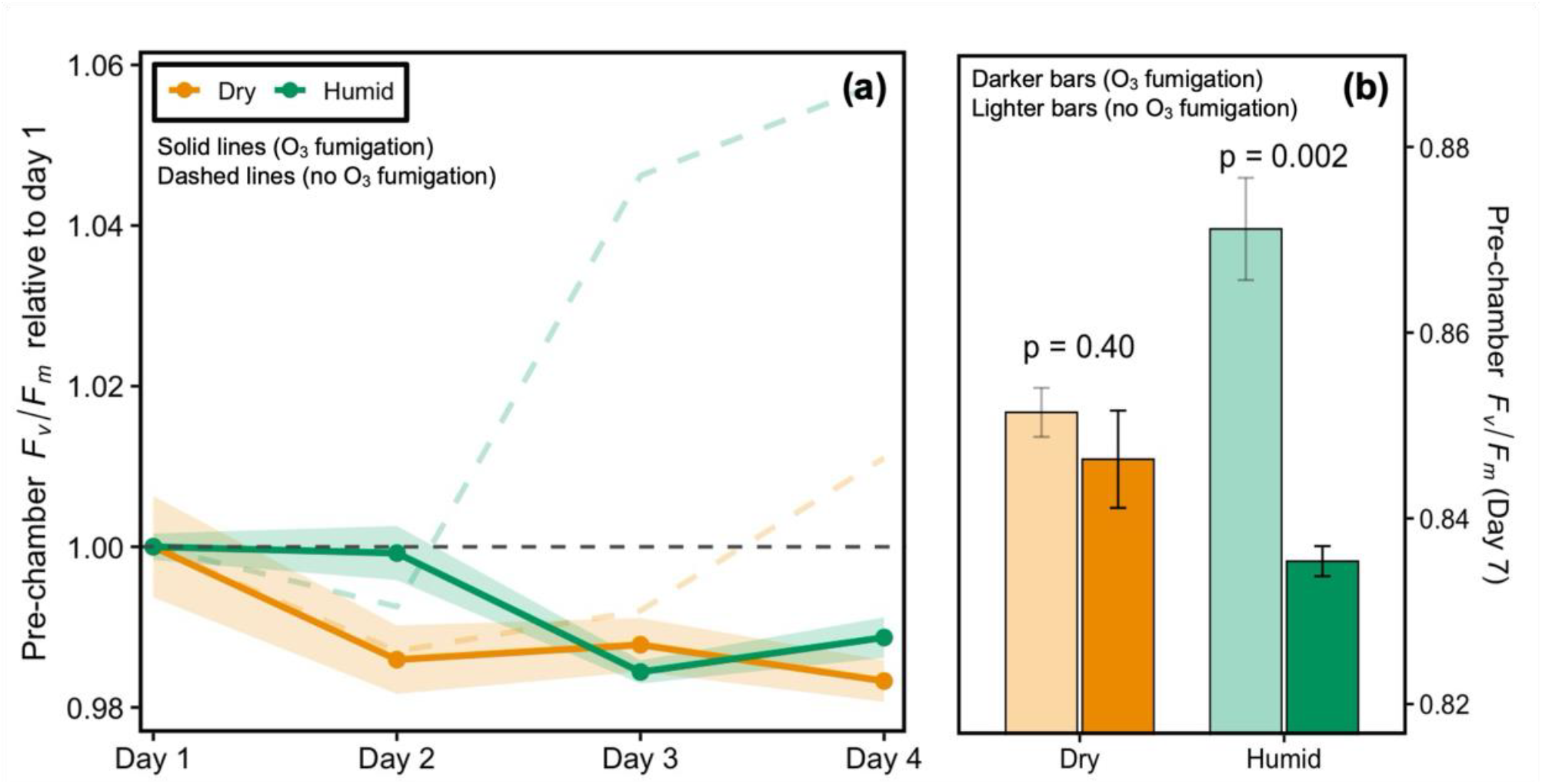
Leaf-level responses of Meyer lemon to repeated O_3_ fumigation under contrasting humidity conditions. (a) Temporal evolution of pre-chamber F_v_/F_m_ during four consecutive exposure days (normalized to the first day pre-chamber value for each treatment). Solid lines and shaded areas represent O_3_-fumigated plants (mean ± SE), while dashed lines show corresponding non-fumigated controls. (b) Mean (± SE) F_v_/F_m_ measured on day 7 (recovery); lighter bars correspond to control plants (no O_3_ fumigation).

Under dry air (∼25% RH), F_v_/F_m_ in O_3_-fumigated plants matched the controls through three days of exposure and differed only on fourth day. This one-day delay in O_3_ defense relative to the humid treatment (**Fig. 1a**) could have been crucial to the observed recovery (**Fig. 1b**), where F_v_/F_m_ had rebounded and was not significantly different from the controls (p = 0.40).

Exposure magnitude was intentionally subtle and realistic (80 ppb, 4 h × 4 days); nevertheless, these modest, episodic exposures produced measurable F_v_/F_m_ impairment within 3-4 exposure days, with a humidity-dependent timing and recovery.

Earlier response to elevated O_3_ in the high humidity can be explained by higher stomatal O_3_ flux. Higher humidity maintains higher g_s_ which accelerates O_3_ entry and reactive oxygen species (ROS) formation in the apoplast and at the thylakoid, which depresses F_v_/F_m_ (Betzelberger et al., 2012; Mills et al., 2007; Tai et al., 2021). This humidity-stomata-dose linkage is a central reason flux-based metrics often outperform concentration metrics in predicting injury (Mills et al., 2007; Tai et al., 2021; Van Dingenen et al., 2009) and is consistent with large-scale observations that VPD is the strongest predictor of midday O_3_ via stomata-regulated dry deposition – high VPD (dry air) restricts stomata, reducing O_3_ uptake, whereas moist air promotes uptake (Kavassalis & Murphy, 2017). Under our humid treatment, stomata likely stayed open across the week, causing the defense to break on the third day and remain depressed after two days without O_3_ exposure (day 7; **Fig. 1b**)

Under dry air, partial stomatal closure likely delayed O_3_ uptake and postponed the O_3_ impact on F_v_/F_m_ until fourth day which was followed by complete recovery by day 7 (**Fig. 1b**) – implying that, at this subtle dose, oxidative damage was transient rather than structurally irreversible. This protective delay under dryness supports crop meta-analyses showing that water stress can reduce O_3_ dose (via g_s_) and postpone injury (Feng & Kobayashi, 2009; Tai et al., 2021). It also aligns with modeling work demonstrating the sensitivity of O_3_ dry-deposition pathways to plant ecophysiology and water status (Sun et al., 2022; Wong et al., 2018)

Observed decoupling from control measurements under dry air on fourth day of exposure (**Fig. 1a)** shows that non-stomatal pathways (cuticle, apoplast chemistry, in-canopy reactions, soil uptake) and cumulative oxidative load still drive injury once exposure continues – thus, dryness did not prevent injury, only delayed it. Recent analyses indicate that non-stomatal O_3_ uptake can increase during hot episodes and, more broadly, that current big-leaf parameterizations often under-capture temperature and dryness responses of non-stomatal deposition (Sun et al., 2022; Wong et al., 2022). Therefore, the observed pattern of slower onset under dry air followed by rapid recovery is something to be expected if stomatal limitation initially reduces dose, while non-stomatal routes deliver O_3_ more slowly, accumulating enough oxidative stress to tip F_v_/F_m_ on the fourth day. It is likely that additional exposure beyond day 4 would have prohibited recovery in dry-air plants, similar to the suppression observed for the humid treatment – but for that, longer experiments are needed.

Although the O_3_ exposure in our experiment was subtle, it is representative of realistic – under human health thresholds (the National Ambient Air Quality Standard is based on an 8-hour average of 70 ppb) – conditions, and supports some literature which argues that repeated, moderate O_3_ peaks observed during heatwaves and wildfire-impacted periods can significantly affect plant function (Mills et al., 2007; Tai et al., 2021; Van Dingenen et al., 2009). In the U.S., recent events (e.g. Canadian wildfires in 2023) produced early-season high O_3_ anomalies (Cooper et al., 2024), and study by Chang et al. (2025) shows continuing complexity in extreme-O_3_ behavior during heat waves. Our results highlight the influence of environmental conditions (RH or VPD) in determining whether a given ambient O_3_ episode crosses a plant’s physiological threshold faster or slower.

To test if the humidity-driven mechanisms observed at the leaf level also hold under real field conditions, we next analyzed satellite-based canopy fluorescence responses across two contrasting citrus regions.

### 3.2. Regional SIF responses to O_3_ episodes in contrasting citrus climates – humidity accelerates O_3_-induced canopy stress, dryness delays it

To evaluate whether the mechanistic differences observed at the leaf level scale up to regional canopies, we analyzed weekly SIF over two major citrus-growing regions that contrast in climate: Fresno County, California (hot semi-arid) and Indian River County, Florida (humid subtropical) (**Fig. 2c**).

**Fig. 2.**
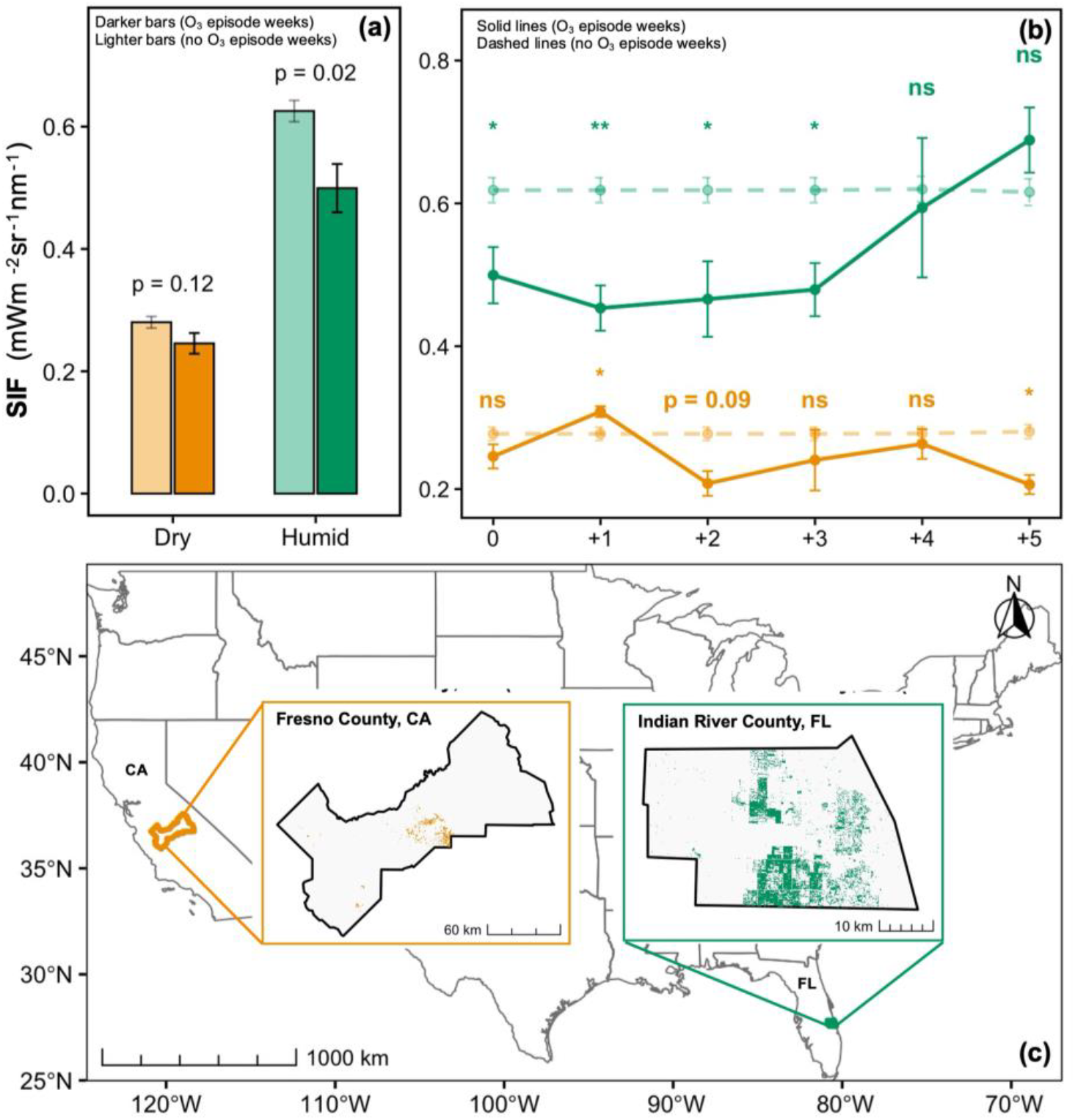
Regional-scale SIF responses to O_3_ episodes in citrus-growing regions with contrasting climates. (a) Mean weekly SIF during O_3_-episode weeks (dark bars) and non-episode weeks (light bars) in Fresno County, CA (dry, hot semi-arid) and Indian River County, FL (humid subtropical). Error bars show ± SE. (b) Temporal evolution of mean SIF for O_3_-episode weeks (solid lines) and non-episode weeks (dashed lines) over five weeks following O_3_-episodes. Markers indicate significance of difference between the two composites for each week (ns = not significant; \**p* < 0.05; \*\**p* < 0.01). (c) Locations of citrus fields analyzed in each region.

Across all weeks, SIF was higher in the humid subtropical region than in the dry semi-arid region (**Fig. 2a**), reflecting higher citrus productivity and longer photosynthetic seasons in humid climates (Leng et al., 2022; Miao et al., 2018; Wu et al., 2024). During high O_3_ weeks, SIF in Florida (humid) declined significantly (p = 0.02), while in California the difference was not significant (p = 0.12).

Temporal composites (**Fig. 2b**) show that in humid climate, SIF remained below non-O_3_ baseline (dashed line) for three weeks (+3) after O_3_ episodes, then recovered sharply by the fourth and fifth week. In the dry climate, SIF briefly increased in the first week, then dropped in second week – similar trend was observed under drought stress in several other studies (Chen et al., 2018; Helm et al., 2020; Xu et al., 2021). These trends mirror chamber results – an immediate O_3_ impact under humid air and a delayed response under dry air. Thus, humid conditions favor stomata-regulated dry deposition and stronger O_3_ – meteorology coupling (Kavassalis & Murphy, 2017). Because drought effects were removed, the immediate SIF loss in Florida reflects the O_3_ flux pathway, not water stress. In contrast, dry air limits g_s_ and delays injury – the same kinetic protection observed in chambers. Yet this protection is temporary: non-stomatal O_3_ uptake, which increases under heat and dryness, can still accumulate and trigger later damage (Sun et al., 2022; Wong et al., 2022).

The SIF increase in the first week after the O_3_ event in dry climate, followed by a drop in the second week, could have two possible explanations supported by the literature. Firstly, under dry conditions, SIF-GPP coupling is driven by physiology, not structure (Wu et al., 2024). A transient relief of VPD or temperature can reopen stomata and enhance electron transport, producing a short-lived SIF rebound (Miao et al., 2018; Wu et al., 2024), while non-stomatal injury continues to build (Sun et al., 2022; Wong et al., 2022). Similar short-term rebounds under water stress are observed when temporary relaxation restores SIF before oxidative damage resurfaces (Chen et al., 2018; Helm et al., 2020; Xu et al., 2021). Secondly, in California, aerosols can co-vary with O_3_ episodes. Particulate matter pollution suppresses photosynthesis (He et al., 2023) and wildfire smoke (common in California) modulates absorbed photosynthetically active radiation and SIF (Chang et al., 2025; Cooper et al., 2024) which may be a reason to different SIF reaction, but further research is needed.

The immediate and persistent SIF suppression in humid citrus orchards confirms that humidity amplifies the effective O_3_ dose received through the stomata promoting rapid production of ROS that down-regulate F_v_/F_m_ and photosynthetic electron transport (Betzelberger et al., 2012; Mills et al., 2007; Tai et al., 2021). The sharp recovery through weeks four and five indicates canopy repair once O_3_ episodes end, consistent with SIF flux tower measurements showing SIF-GPP recovery after drought episodes (Leng et al., 2022; Miao et al., 2018; Wu et al., 2024). Flux-based risk frameworks similarly predict stronger and longer responses to O_3_ exposure under humid air (Mills et al., 2007; Tai et al., 2021).

Evergreen citrus canopies maintain photosynthetically active foliage through O_3_ seasons, so even moderate, but repeated O_3_ peaks can impact photosynthetic function without visible injury, which SIF captures in both climates (Ainsworth et al., 2014; Montes et al., 2022). The humid region shows immediate and sustained losses, while the dry region exhibits a delayed response with a short rebound. Recent U.S. records of wildfire-driven O_3_ anomalies (Chang et al., 2025; Cooper et al., 2024) reinforce that these episodic dynamics are realistic. Because stomatal behavior depends on humidity/VPD, the same AOT40 exposure yields different effective doses in Florida and California – exactly what our SIF composites reveal (Kavassalis & Murphy, 2017; Sun et al., 2022)

Overall, humidity accelerates and prolongs O_3_ damage, while dryness delays it but cannot prevent it. Non-stomatal pathways ultimately impose a slower, lingering stress. This continuum – from fast stomatal injury in humid air to delayed non-stomatal damage in dry air – links our chamber results to real orchards and underscores SIF’s value as an early-warning signal of subtle O_3_ stress in citrus orchards.

### 3.3. SIF-GPP relationship under drought and O_3_ stress – dominant stressor shifts with climate

Because SIF reflects light-use efficiency while GPP integrates carbon assimilation, their coupling provides a direct link between canopy photochemistry and productivity under stress (Miao et al., 2018; Wu et al., 2024). Relationships between SIF and GPP differed across climates and stress types (**Fig. 3**). In the dry semi-arid region, the SIF-GPP link remained stable during drought weeks, supporting that citrus orchards are already physiologically adjusted to periodic water stress (Leng et al., 2022; Wu et al., 2024). Perennials such as citrus likely maintain photosynthetic coordination under moderate dryness through sustained photoprotective control and stable canopy structure (Helm et al., 2020; Miao et al., 2018). However, during high-O_3_ weeks the SIF-GPP relationship collapsed, indicating that O_3_ stress disrupts electron transport and carbon assimilation simultaneously (Betzelberger et al., 2012; Mills et al., 2007; Tai et al., 2021), breaking the usual linear coupling between fluorescence and productivity (Mamić et al., 2025b; Pickering et al., 2022) – supporting our earlier observation that, in dry air, stomatal limitation delays but does not prevent oxidative injury and that non-stomatal O_3_ uptake eventually drives canopy-level dysfunction (Sun et al., 2022; Wong et al., 2022).

**Fig. 3.**
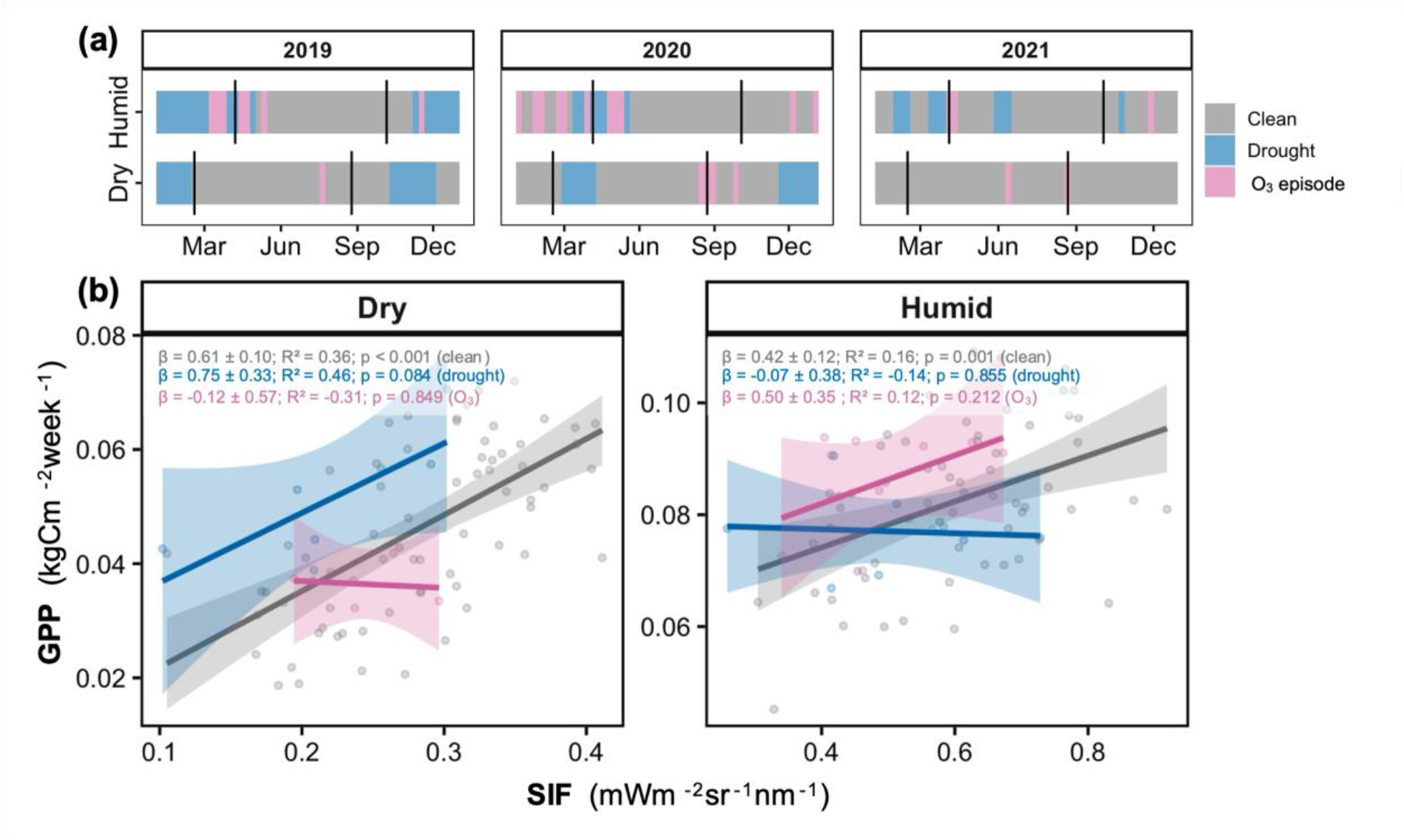
Relationships between SIF and GPP in citrus regions under clean, drought, and O_3_ conditions. (a) Temporal distribution of analyzed periods (marked with vertical black lines) from 2019-2021 showing clean (gray), drought (blue), and O_3_ (pink) weeks over citrus fields in Fresno County, CA (dry) and Indian River County, FL (humid). (b) Linear regressions between SIF and GPP for each condition and climate during analyzed periods. Shaded areas represent 95% confidence intervals for fitted slopes (β ± SE).

In the humid subtropical region, the trend was reversed. Here, drought weeks led to a weaker or fully decoupled SIF-GPP relationship, while O_3_ weeks were similar to clean conditions. This suggests that citrus in humid climates may be more vulnerable to episodic dryness than to O_3_, likely because trees acclimated to high humidity maintain high g_s_ and are less prepared for rapid water limitation (Ainsworth et al., 2014; Montes et al., 2022). The relatively stable SIF-GPP coupling during O_3_ weeks could reflect coordinated stomatal regulation, as higher humidity maintains g_s_ and synchronizes O_3_ flux with CO2 exchange (Mills et al., 2007; Tai et al., 2021).

Rather than a direct comparison with the week-by-week SIF analysis in section 3.2, these regressions identify which limitation dominates under each climate. In the humid region, drought induces stomatal closure and high non-photochemical quenching (NPQ), reducing carbon assimilation more than fluorescence and thus eroding the SIF-GPP link (Helm et al., 2020; Miao et al., 2018; Wu et al., 2024). In the dry region, drought responses are largely acclimated, preserving coupling, whereas O_3_ exposure causes photochemical and oxidative damage, including delayed non-stomatal injury, that ultimately breaks proportionality between SIF and GPP (Sun et al., 2022; Wong et al., 2022).

While interpreting these regressions, we note that modelled MODIS GPP carries an uncertainty of approximately 20-30% (Running et al., 2004; Wang et al., 2017). Therefore, we focus on relative patterns between regions which suggest that the dominant stressor controlling canopy photosynthetic coupling shifts with climate – O_3_ dominates under dry air, while drought dominates under humid air.

## 4. Conclusions

This study demonstrates that atmospheric humidity controls how citrus responds to O_3_ stress across scales – from leaf-level F_v_/F_m_ to canopy-level SIF and GPP. Under humid air, high g_s_ accelerates O_3_ uptake, leading to early and persistent declines in F_v_/F_m_ and SIF, while under dry air, stomatal limitation delays but does not prevent oxidative injury. The same mechanisms extend to the regional scale where O_3_ dominates canopy decoupling in dry climate, whereas drought dominates in humid climate.

These findings align with and extend previous citrus and crop O_3_ studies (Ainsworth et al., 2014; Calatayud et al., 2006; Delgado-Saborit & Esteve-Cano, 2008; Fares et al., 2010; Montes et al., 2022). Earlier work showed that O_3_ reduces citrus photosynthesis and pigment content primarily through stomatal uptake, while Fares et al. (2010) emphasized the tight coupling between citrus physiology, biogenic volatile organic compounds, and local O_3_ chemistry. Our results add that the humidity context determines when this physiological threshold is crossed and whether canopies recover. The identified humidity-dependent kinetics, i.e. response reaction speed, help reconcile inconsistencies among field observations of citrus O_3_ sensitivity across climates and provide a mechanistic basis for interpreting satellite SIF signals in perennial crops. Because citrus orchards maintain evergreen foliage through the O_3_ season, even moderate, but repeated episodes can impose cumulative strain detectable well before visible injury – an effect likely underrepresented in global yield models (Leung et al., 2022; Tai et al., 2021).

In practical terms, this study highlights that O_3_-risk assessments must integrate local humidity and VPD conditions to accurately capture effective O_3_ dose. Incorporating both stomatal and non-stomatal deposition pathways into regional models will improve projections of O_3_ impacts on citrus productivity under future climate and air-quality scenarios. Remote sensing data, such as TROPOMI SIF, provide a scalable tool for monitoring these subtle stresses in real time. The main limitation of TROPOMI is its coarse spatial resolution of 3.5 × 7 km. However, the future ESA FLEX mission will provide global SIF measurements with a spatial resolution of 300 m, opening the possibility of scaling these methods down to a smaller field scale.

## Acknowledgments

This paper and related research have been conducted within the framework of the Italian National PhD Program in Earth Observation funded by the European Union – Next Generation EU through the Project of Italian Recovery and Resilience Plan (NRRP) (B53C22004370006). The TROPOSIF products were generated by the TROPOSIF team conducted by NOVELTIS under the European Space Agency (ESA) Sentinel-5p+ Innovation activity Contract Nº 4000127461/19/I-NS. We acknowledge funding from a National Science Foundation AGS Postdoctoral Research Fellowship (MR; 2306215). We would like to acknowledge the help of Rose Rossell and Tony Hernandez, who not only assisted in setting up our chamber experiment, but also took PPS measurements when we were unable to do so.

