## Supplementary material for "Humidity Controls the Timing and Persistence of Ozone Injury in Citrus: Linking Leaf Physiology and Regional Canopy Responses": Fig. S1; Fig. S2

**Supplementary Information**

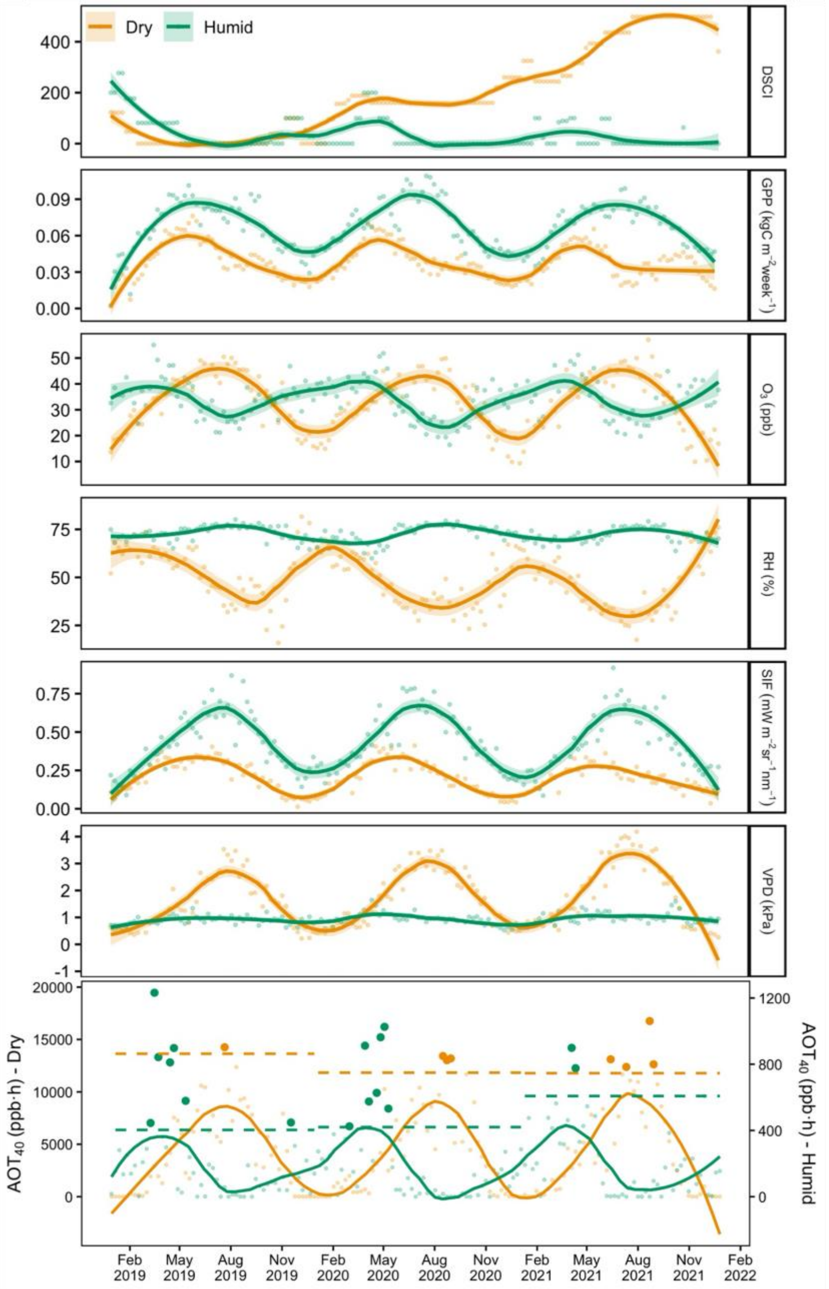


**Figure S1.** Weekly time series of DSCI, GPP, $O_3$, RH, SIF, VPD, and AOT40 for citrus orchards in a dry (Fresno County, CA; orange) and humid (Indian River County, FL; green) region during 2019-2021. Solid lines show LOESS fits (95 % confidence intervals) of weekly means. Dashed lines indicate region-specific 3 × median absolute deviation (MAD) thresholds used to define $O_3$ episodes, and opaque points mark weeks exceeding those thresholds. The dry region shows higher $O_3$ accumulation, VPD, and DSCI but lower RH, SIF, and GPP than the humid region, reflecting distinct climatic and physiological regimes.

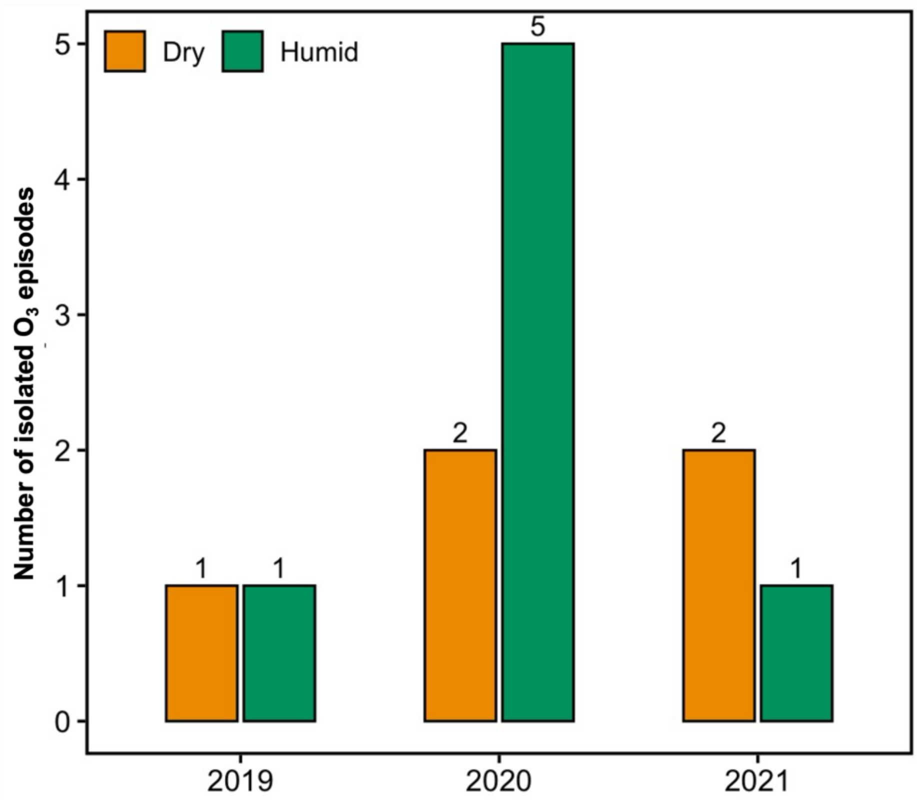


**Figure S2.** Counts of isolated high-$O_3$ episodes (weeks) by year in the two study regions: Dry (Fresno County, CA; orange) and Humid (Indian River County, FL; green).
